# ABTrans: A Transformer-based model for predicting interaction between anti-Aß antibodies and peptides

**DOI:** 10.1101/2023.10.09.561503

**Authors:** Yuhong Su, Lingfeng Zhang, Yangjing Wang, Buyong Ma

## Abstract

Understanding the recognition of antibodies and Aβ peptide is crucial for the development of more effective therapeutic agents. Here we studied the interaction between anti-Aβ antibodies and different peptides by building a deep learning model, using the dodecapeptide sequences elucidated from phage display and known anti-Aβ antibody sequences collected from public sources. Our multi-classification model, ABTrans was trained to determine the four levels of binding ability between anti-Aβ antibody and dodecapeptide: not binding, weak binding, medium binding, and strong binding. The accuracy of our model reached 0.8278. Using the ABTrans, we examined the cross-reaction of anti-Aβ antibodies with other human amyloidogenic proteins, and we found that Aducanumab and Donanemab have the least cross-reactions with other human amyloidogenic proteins. We also systematically screened all human proteins interaction with eleven selected anti-Aβ antibodies to identify possible peptide fragments that could be an off-target candidate.

**Key Points:**

- ABTrans is a Transformer-based model that was developed for the first time to predict the interaction between anti-Aß antibodies and peptides.
- ABTrans was trained using a dataset with 1.5 million peptides and 110 anti-Aβ antibodies.
- ABTrans achieved an accuracy of 0.8278 and is capable of determining the four levels of binding ability between antibody and Aß: not binding, weak binding, medium binding, and strong binding.
- ABTrans has potential applications in predicting off-target and cross-reactivity effects of antibodies and in designing new anti-Aß antibodies.

## Introduction

Alzheimer’s disease (AD), one of the most common forms of dementia[1], is a neurodegenerative disease characterized by memory loss, cognitive impairment, and personality change. It has become a worldwide public health dilemma and has a significant impact on the direct cost of AD to society[2–4]. It is expected that by 2050 there will be 115 million AD patients. The typical pathological sign of AD is a large number of senile plaques (SP) and the accumulation of neurofibrillary tangles (NFTs) in the patient’s brain. β amyloid protein(Aβ) is the core component of senile plaques and the NFTs is formed by the misfolding and aggregation of abnormal hyperphosphorylated tau proteins. Both Aβ and tau protein play key roles in the occurrence and development of AD[5]. Aβ is generally composed of 40-42 amino acids and is the enzymatic hydrolysis product of amyloid precursor protein (APP), in which Aβ1-42 are more likely to aggregate and form amyloid deposits. The aggregates of Aβ have obvious direct or indirect toxic effects on nerve cells, leading to the reduction of neural synapses and neuronal necrosis [6, 7]. Aβ aggregates include dimers, oligomers, and fibrils according to their sizes and structures,[8] while oligomeric Aβ are reported to have stronger cytotoxicity.[9–11] Therefore, clearing aggregated Aβ, especially oligomers, has emerged as the major therapeutic goal. The candidates of anti-Aβ drug [12, 13] include nanoparticles,[14] peptides,[15] small molecules[16, 17] and antibodies.[18] While using antibodies to reduce assembled Aβ provide a promising approach and have been widely investigated,[18] many antibodies failed in various clinical stages, such as bapineuzumab[19], solanezumab [20], and crenezumab[21]. The difficulties in amyloid antibodies therapies have triggered debates about amyloid hypothesis.[22, 23] Nevertheless, recently FDA approvals of aducanumab and lecanemab provided important validations of amyloid as a therapeutic target. [24–29]

Amyloids are known for their polymorphic conformations, [30] and Aβ forms multiple structures of oligomers/fibrils under different conditions. [31–36] Different soluble Aβ aggregates can give rise to cellular toxicity through different mechanisms.[37] It is very hard to control the selectivity of amyloid-directed antibodies for the soluble Aβ oligomers. [38] Furthermore, some amyloid antibodies can recognize amyloids with similar conformations but with different amino acid sequences, for example different Aβ mutants [39] or totally different proteins. [40] Therefore, it is very important to understand an antibody’s sensitivity against different amyloidogenic proteins.

Previously we have used structural analysis[41] and molecular dynamics simulations[42–45] to gain insights into specific antibodies’ recognition of amyloid monomers, oligomers, and fibrils. In order to catch general features of antibodies’ recognition of Aβ-like sequences and implicated conformations, we developed a deep learning model ABTrans to predict the interaction between antibodies and dodecapeptides. To our knowledge, it is the first time that deep learning was used in the study of passive immunotherapy for Alzheimer’s disease, and different from the general interaction prediction model, which only can predict whether it binds[46].

ABTrans can predict the four binding modes: strong, medium, weak and non-binding. Based on the large number of dodecapeptide data obtained by phage display technology and the antibody sequences collected from the public resources, a deep learning model based on the transformer framework was constructed to predict the binding of dodecapeptide and antibody. The accuracy of our model reached 0.8278. After that, the model is applied to study the off-target and cross-reactivity effects of antibodies and found that aducanumab may have the least off-target proteins and better specificity.

## Materials and Methods

### 1. Data acquisition and preprocessing

Previously, a phage display experiment was conducted to screen the dodecapeptides that bind anti-Aβ antibodies [47]. Twenty-eight monoclonal antibodies were used to pan 10^11^ phage from a dodecapeptide library with a complexity of 10^9^ sequences. Three rounds of panning and amplification were carried out. Phages obtained at each step were sequenced to analyze. We used the peptide sequences to construct our dataset. Each round’s data are labelled ‘Positive Low’, ‘Positive Intermediate’, ‘ Positive’, and ‘Positive High’. The polypeptide dataset is de-duplicated. The sequences in each round that are similar to other rounds with more than 70% identity are removed by the CD-HIT program[48] and saved in the round with higher affinity. The ‘Negative’ dataset (noninteraction pairs) is randomly generated dodecapeptides, which are further screened by removing sequences sharing more than 60% sequence identity with other groups.

Antibodies that bind Aß are collected from published literature and patents. the full sequence is encoded with the Chothia numbering scheme[49], hence every amino acid has its own number. Since sometimes the two residues at both ends of a CDR sequence are found to engage in binding[50], we use each CDR, as well as the additional four external residues, as an independent training sequence. The total sum of CDR sequences is designed to be 101 amino acids, large enough to include most known anti-Aß antibodies. To maintain the precise position of amino acids, the gap positions within 101 amino acid input window were labeled as letter “U”.

The numbers of peptides in the four data groups are different (Supplementary Table 1), so we under-sampled the peptide number to contain 267884 samples in each group (Supplementary Table 2). 110 antibodies are paired with each category (random or exhaustively). Finally, the whole dataset was divided into the training set, validation set and testing set according to the ratio of 70:15:15.

### 2. Model

The ABTrans framework learns to predict the interaction of Aβ peptides with antibodies as follows (Fig. 1b). First, both antibody and antigen (peptide) are encoded by the sum of input embedding and positional embedding. Then, they are processed by two different encoder modules respectively. The encoder layer contains the masked multi-head self-attention, layer normalization layer, and position-wise feed-forward layer. For the purpose of fusing information of both antibody and antigen, the outputs of encoders are concatenated as the embedding of the next encoder. At last, a stacked fully connected module is used to increase the nonlinear expression ability of the model and predict the classification of their affinity.

**Figure 1.**
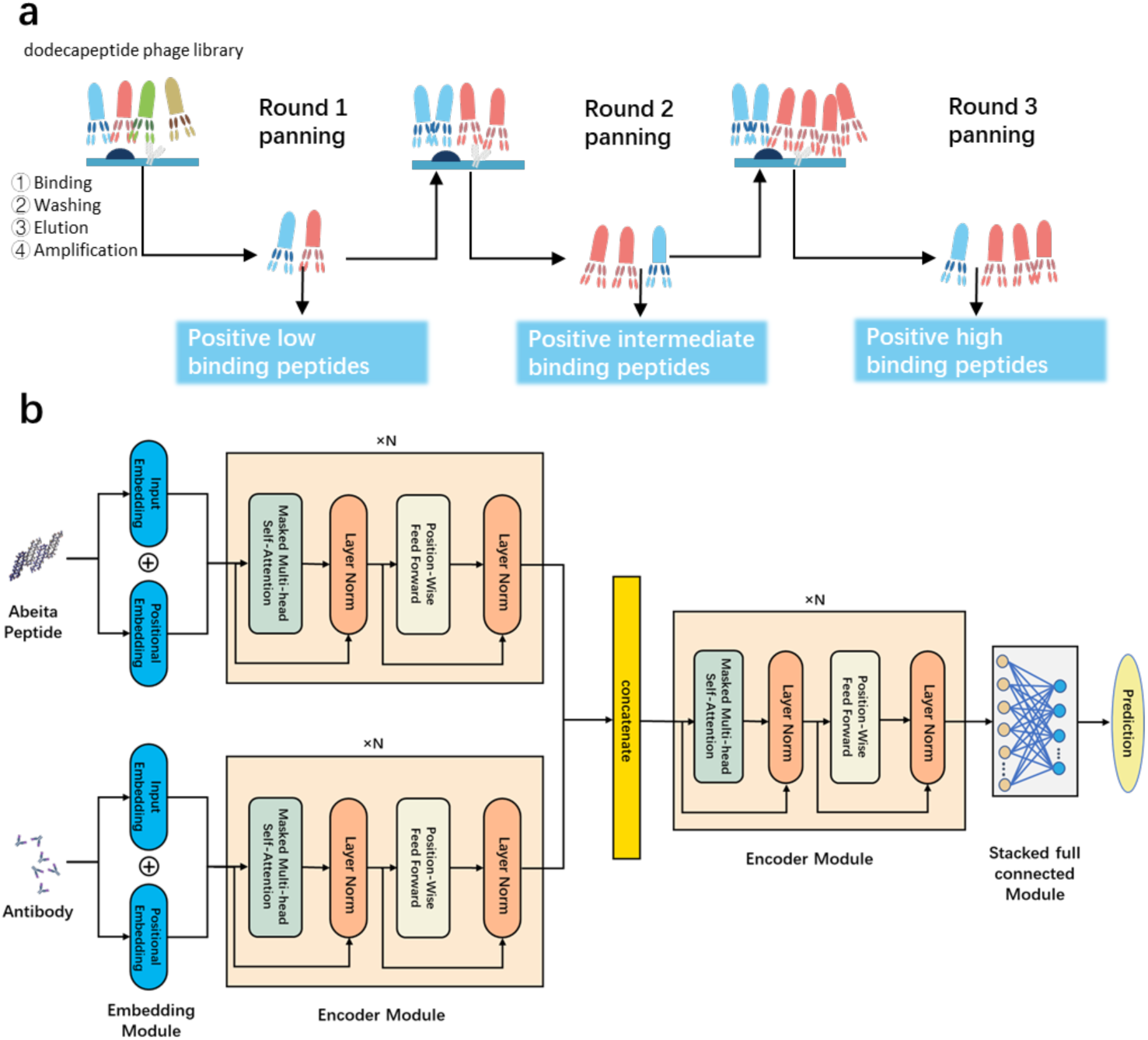
deep learning model for predicting the interaction of anti-Aβ antibodies and dodecapeptide. (a) Experimental workflow for generation dodecapeptide. Three rounds of panning and amplification were carried out[47]. (b) Transformer model framework for predicting the binding of antibodies against Aβ to dodecapeptide.

#### 2.1 Sequence embedding

First of all, each amino acid is represented by a unique number. Then, the Aβ peptide and antibody sequences are constructed by these numbers. Since this amino acid number is too simple, an embedding layer is devised, which transforms the number to continuous values, which may capture the relationship between amino acids in a high-dimensional space, which is also called the latent embedding space.[51]

The next step is to add positional information into the embeddings since the order of sequence is essential for both amyloid beta peptide and antibody. We use this formula to calculate:

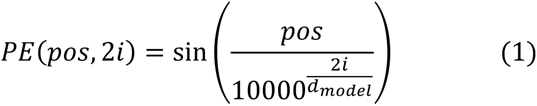

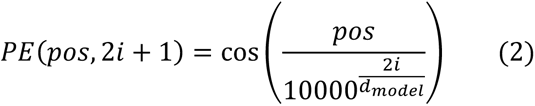

Where *pos* is the position of amino acid in the sequence, *d_model_* represents the input vector’s dimension and *i* is the *i*th element of this vector. For every odd index on the input vector, a positional encoding is created using the cosine function. For every even index, a positional encoding is created using the sine function. This positional encoding not only can capture the absolute position but also the relative position information in the sequence[52]. At last, those positional encodings are added to their corresponding input embeddings. In Supplementary Fig. 1, we illustrated the embeddings of a peptide with 12 different amino acids.

#### 2.2 Masked mechanism

The self-attention mechanism enables the transformer to extract the long-term dependency [52]. Inspired by a transformer-based model to predict peptide–HLA class I binding[53], the masked multi-head self-attention mechanism is devised as the core of the encoder in the ABTrans model. It applies a specific attention mechanism called self-attention.

Given an input X, where X can be either an Aβ peptide or an antibody sequence, it will be fed into three linear layers to get Q(query), K(key), and V(value) matrix.

Then, an attention score matrix is generated by a scaled dot product from the Q and K matrix. Here, the attention score determines the degree of importance or relevance of each amino acid in the input sequence to the current amino acid being processed.

After that, the V matrix and the attention score matrix are multiplied to get new latent amino acid embeddings. Self-attention expands the model’s ability to focus on different positions and multi-head allows the model to create multiple “representation subspaces” in different ways. For the purpose of masking out the last padded “fake” amino acid of the output sequence, -1e9 is used instead of these actual values.[52]

Multi-head self-attention can be calculated by following formulas.

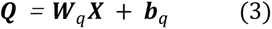

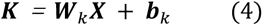

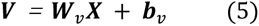

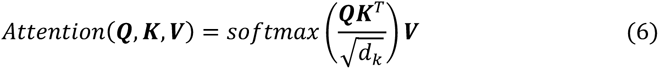

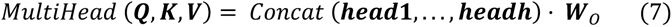

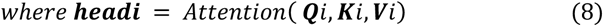

where *d_k_* refers to the dimension of ***K*** vector, ***Q***/***K***/***V*** represents the Query/Key/Value vector. ***W**_q_*, ***b**_q_*, ***W**_k_*, ***b**_k_*, ***W**_v_*, ***b**_v_* are the three different sets of linear parameters.

#### 2.3 Hyper-parameter setting

The dimension of the word vector corresponding to an amino acid is mapped to 64, the dimension of the transformation matrix in the feedforward fully connected network to 512, and the dimension of each head word vector of multi-head self-attention to 64. On account of the limited computing resources, we focused on adjusting the number of layers of the encoder and the number of heads in the multi-head self-attention mechanism.

#### 2.4 Implementation

The models were trained on the High-Performance Computing of Shanghai Jiao Tong University with two A100 GPUs which have 80G memory in total. The CPU is Intel Xeon ICX Platinum 8358 @ 2.60GHz. The models were built by PyTorch 1.9.0 and Python 3.6.13. During training, the maximum epoch is set to 50. The batch size is 128 for each GPU. The initial learning rate is set to 1e-3. If the training accuracy is no longer improved, the learning rate will be halved. If the loss doesn’t decrease five times, the model will stop training[54]. We use the Adam optimizer[55] and cross-entropy loss function in our model.

### 3. Performance metrics

For each model, the following metrics were calculated:

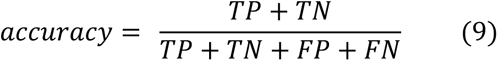

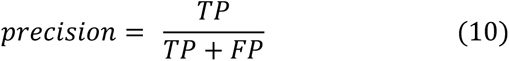

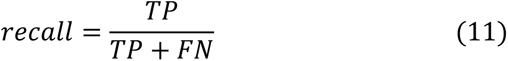

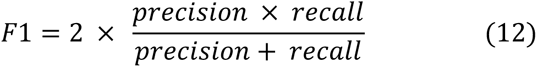

where TP, TN, FP, and FN are the numbers of true positives, true negatives, false positives, and false negatives, respectively. In addition, the area under the receiver operating characteristic curve (AUC) and the area under the precision-recall curve (AUPR) are adopted as the other performance evaluation metrics. The higher the value of the metric, the better the model. The confusion matrix also is drawn to see the distribution of each sample.

### 4 The off-target effects of antibody

23391 homo sapiens protein structures are downloaded from the AlphaFold database[56], and their sequences are extracted. Then, every protein is divided into dodecapeptides in the origin sequence order. The antibodies include seven drug antibodies (Bapineuzumab, Solanezumab, Crenezumab, Ponezumab, Gantenerumab, Aducanumab, and Donanemab[57–63]) and four antibodies (mA11-204, mA11-205, mA11-118, and mA11-55) from the experiment[47]. To analyze the antibody’s off-target effects, the best model is applied to predict the interaction between peptides belonging to human protein and antibodies.

According to the test result, the peptides interacting strongly with the antibody will be analyzed further. The python script is used to filter these peptides by removing two Protine, all residues are hydrophobic or none of them have charged group. With the intention of knowing whether these peptides are on the surface of the protein, the rest of the peptides’ solvent-accessible surface area (SASA) is calculated by freesasa program[64]. Then, the SASA of peptide below 65% of the maximum value will be removed.

## Result

### Optimization and evaluation of ABTrans

With the increase of panning steps in the phages display, the antigen antibody affinity usually increases. Therefore, we defined the peptides identified in the first round of panning and amplification as weakly binding group, those from the second round as moderately binding, and the third round as the most strongly binding group. At the same time, anti-Aβ antibody sequences are obtained manually from literature and patents to the best of our ability. As a result, 110 unique sequences were found (Supplementary information2).

A deep learning model called ABTrans was constructed to study the interaction between anti-Aß antibody and dodecapeptide. As mentioned previously, the model aims to predict the binding of the antibody and the dodecapeptide with four types: strong, medium, weak, and non-binding. The antibodies were paired with the peptides in each group. The transformer model architecture is a widely used model with good effects in most cases[65]. Compared with convolutional neural networks (CNNs), which extract local information commendably, transformer extracts more global information. Compared with recurrent neural networks (RNNs), the transformer realizes parallelization and solves the long-term dependencies problem, so it can process the data faster than RNNs. In addition, Transformer reduces the risk of vanishing and exploding gradients, which happen often in RNNs.[52] Through training, ABTrans is able to predict the different binding capacities of the antibody and dodecapeptide. First, we adjusted the parameters of the model. In view of limited computing resources, model selection was devised on the number of layer and head number of the multi-head attention mechanism. Table 1 illustrates that when the model has two layers and two heads of attention, the accuracy is the highest, reaching 0.8302. However, for the sake of the interpretability of the model to assign amino acids’ contribution, we chose one layer and ten heads as our final model, and its accuracy reached 0.8278, only slightly lower than the best model. It can be seen from Figs. 2a,b that loss continued to decrease and accuracy steadily increased during the training process. Using the early stop strategy[54], ABTrans only needs about 20 epoch to converge, and the speed of each epoch is very fast. It only needs less than 3 min to complete an epoch on a dataset of about a million data. The total training time is less than one hour, which proves the superiority of our model. The areas under the receiver operating characteristic curve and the precision–recall-curve were 0.9668 and 0.9136, respectively, which proved that it could reliably predict the different binding abilities of dodecapeptide and anti-Aß antibody (Fig. 2c, d). From the confusion matrix, we can see that the model could basically distinguish dodecapeptides with different binding capacities, but it seems relatively easy to confuse dodecapeptides with medium binding and strong binding capacity in ABTrans (Fig. 2e).

**Figure 2.**
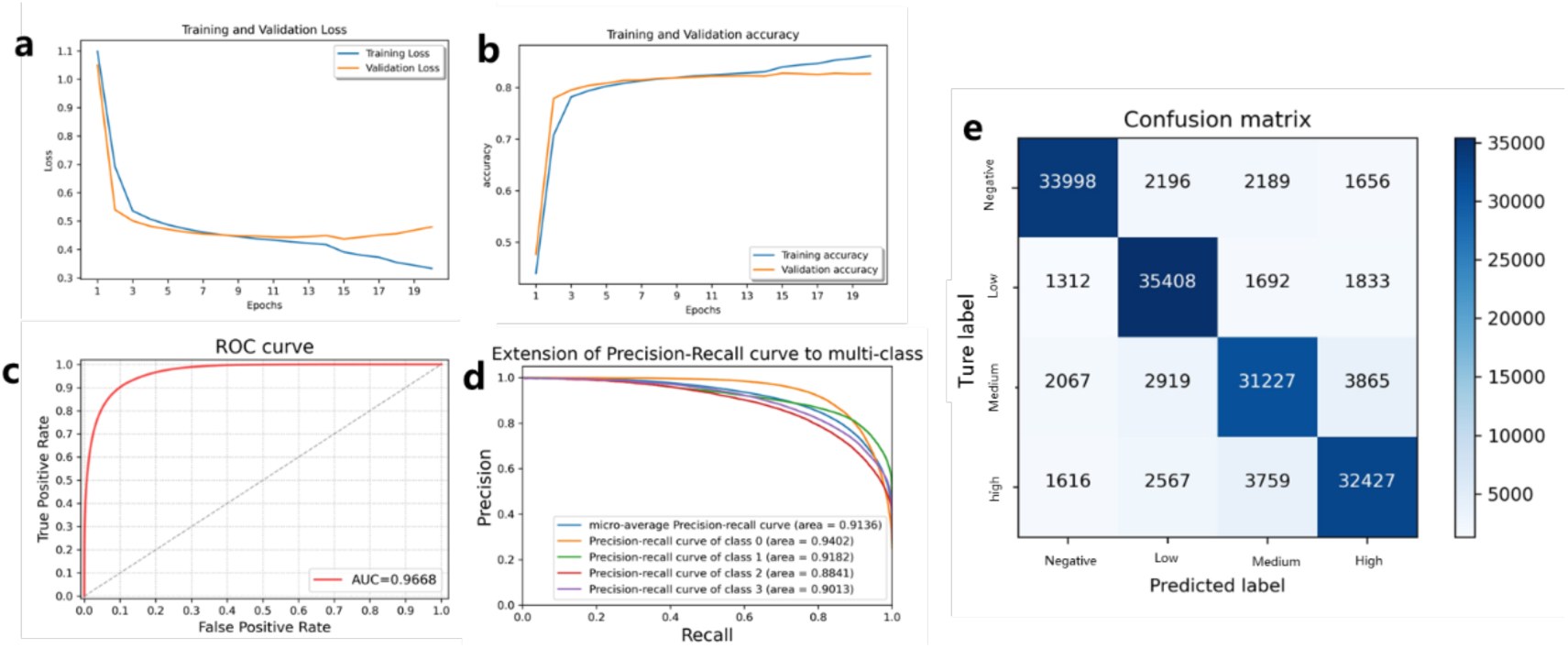
Transformer final model training and test results (a) training and test loss change (b) training and test accuracy change (c) receiver operating characteristic (ROC) curve. AUC, the area under the curve. (d) the precision–recall-curve (e) confusion matrix, showing the number of strong, medium, weak, and non-binding sequences of dodecapeptide

**Table 1.** The metric scores under different hyper-parameters. (where 1-1 represents the number of encoder layers - the number of multi heads self-attention)

|  | ACC | AUPR | AUC | Precision | Recall | F1 |
| --- | --- | --- | --- | --- | --- | --- |
| 1-1 | 0.8196 | 0.9062 | 0.9639 | 0.8198 | 0.8196 | 0.8193 |
| 1-2 | 0.8244 | 0.9106 | 0.9656 | 0.8245 | 0.8244 | 0.8241 |
| 1-3 | 0.8254 | 0.9114 | 0.9659 | 0.8256 | 0.8254 | 0.8251 |
| 1-4 | 0.8252 | 0.9113 | 0.9659 | 0.8253 | 0.8252 | 0.8248 |
| 1-5 | 0.8273 | 0.9135 | 0.9668 | 0.8273 | 0.8273 | 0.8269 |
| 1-6 | 0.8254 | 0.9116 | 0.9660 | 0.8262 | 0.8254 | 0.8255 |
| 1-7 | 0.8224 | 0.9107 | 0.9656 | 0.8234 | 0.8224 | 0.8224 |
| 1-8 | 0.8272 | 0.9126 | 0.9664 | 0.8277 | 0.8272 | 0.8270 |
| 1-9 | 0.8256 | 0.9131 | 0.9666 | 0.8256 | 0.8256 | 0.8253 |
| 1-10 | **0.8278** | **0.9136** | **0.9668** | **0.8280** | **0.8278** | **0.8276** |
| 2-1 | 0.8299 | 0.9151 | 0.9674 | 0.8308 | 0.8299 | 0.8299 |
| 2-2 | 0.8302 | 0.9148 | 0.9672 | 0.8301 | 0.8302 | 0.8296 |
| 2-3 | 0.8257 | 0.9125 | 0.9663 | 0.8265 | 0.8257 | 0.8255 |
| 2-4 | 0.8209 | 0.9085 | 0.9646 | 0.8218 | 0.8209 | 0.8205 |
| 2-5 | 0.8140 | 0.9025 | 0.9622 | 0.8151 | 0.8140 | 0.8139 |
| 2-6 | 0.8199 | 0.9085 | 0.9643 | 0.8217 | 0.8199 | 0.8195 |
| 2-7 | 0.4823 | 0.5850 | 0.7848 | 0.4687 | 0.4823 | 0.4690 |
| 2-8 | 0.8238 | 0.9099 | 0.9652 | 0.8249 | 0.8238 | 0.8237 |
| 2-9 | 0.2491 | 0.2496 | 0.4991 | 0.0621 | 0.2491 | 0.0994 |
| 2-10 | 0.4749 | 0.5773 | 0.7799 | 0.4710 | 0.4749 | 0.4546 |

### ABTrans reveals the distributions of antibody binding epitopes of Aß

The transformer’s attention mechanism can provide insights into Aß antibody’s recognition of peptide. Figure 3a indicates that various multi-heads self-attention at the different positions in a dodecapeptide. The first two amino acids in a dodecapeptide are the most important residues, positions 4, 5, 7, and 8 are also decisive. Then, the binding contribution of amino acid types on dodecapeptides with different binding capacities at different positions is analyzed (Fig. 3b). The binding of the dodecapeptide to the antibody is affected by different amino acids of the peptide. For negative samples, the model mainly focuses on the unknown amino acid X, which may be inferred that the polypeptide obtained in the experiment contains less X amino acid. If the first position is Alanine (A), it is more likely to be a positive sample. For positive samples, the model mainly focuses on Aspartate (D), Leucine (L), and Proline (P) at the N-terminus of the dodecapeptide. Most anti-Aß antibodies bind to the N-terminal of amyloid beta in the phage display experiment[47]. Since the first amino acid at the N-terminal is aspartic acid, it seems that ABTrans could recognize this amino acid to predict the binding type. Proline could be used to distinguish between negative and positive samples, since proline is a beta-strand breaker and usually not involved in amyloid formation. The avoidance of proline in Aß antibody’s recognition of peptide clearly reflects that most Aß antibodies bind to Aß oligomer and fibril which will be broken by a proline residue.

**Figure 3.**
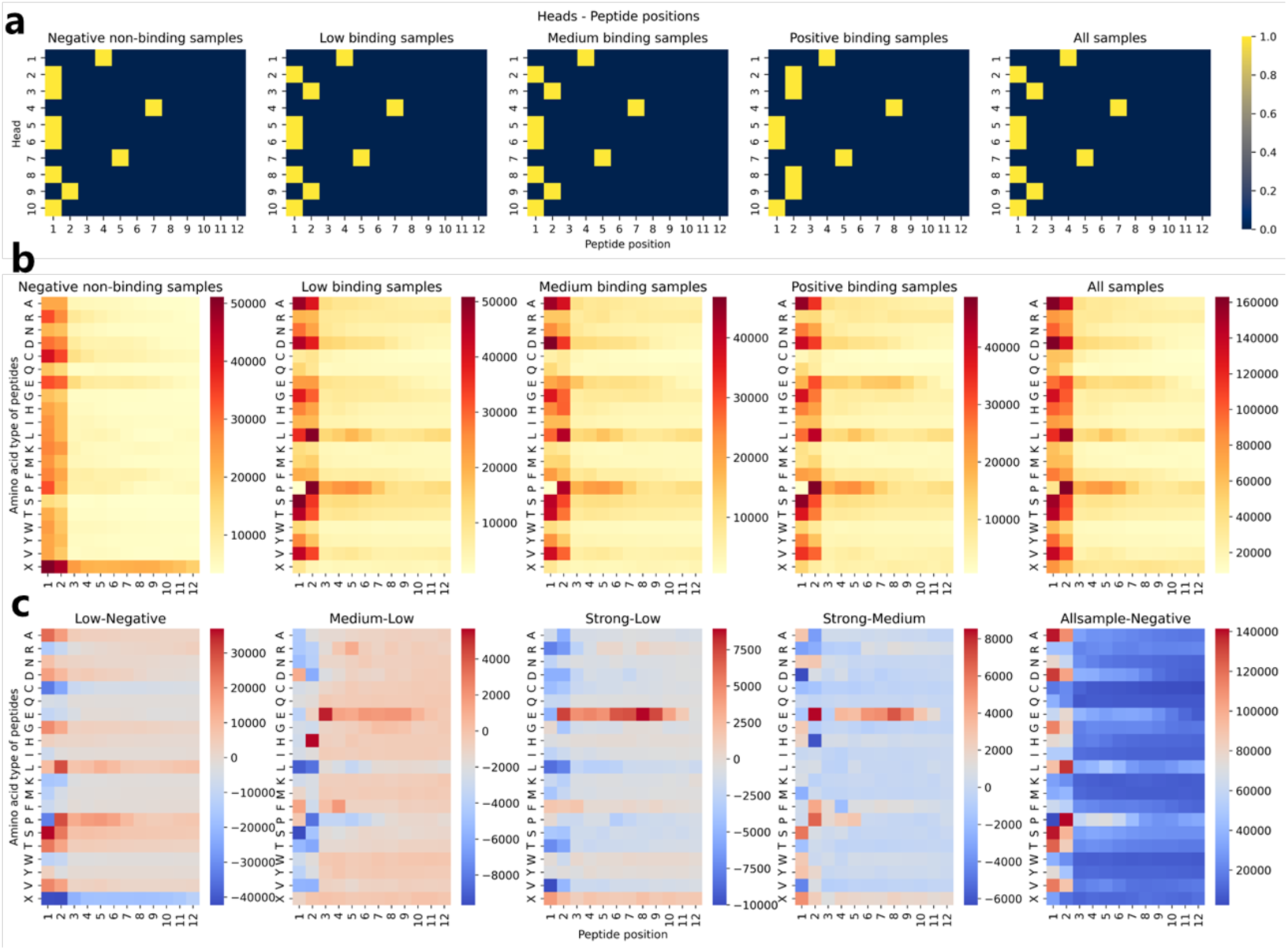
(a) different multi-head self-attention pays attention to different amino acid positions in dodecapeptide. (b) Contribution of amino acid type and position of dodecapeptide to antibody binding (that is, cumulative attention score). (c) The differences between different samples.

From figure 3c, we can find that recognition of glutamic acid is stronger at higher affinity levels. The difference between medium and weak binding lies in the greater emphasis on the presence of glutamic acid at position 3 and histidine at position 2. In comparison to weak binding, strong binding is especially sensitive to the presence of glutamic acid at positions 7, 8, and 9, particularly at position 8. In contrast to medium binding, strong binding is more sensitive to the presence of glutamic acid and proline at position 2, and less sensitive to aspartic acid at position 1 and histidine at position 2.

In general, the attention mechanism can provide insights into residues that were important not only for model classification but also for antigen binding specificity.

### ABTrans prediction of antibody Cross-reactivity effects of by ABTrans

The interaction of Amyloid-beta precursor protein (APP) with the antibody was analyzed to demonstrate the validity of the model. APP is a precursor protein of Aß, and the amino acids at positions 672-713 are Aß sequence. According to the data presented in Table 3, it is evident that all 11 antibodies exhibited robust interactions with the dodecapeptide of amyloid-β. Furthermore, our findings indicate a notable concentration of interacting fragments within the Aβ sequence of the APP protein, indicating that ABTrans presumably could learn the interaction of amyloid and antibody. This also proves the reliability of using this method to study the off-target effects of antibodies.

**Table 2.** Statistics of continuous strong interactions between proteins and antibodies.

| Number | mA<br>11-55 | mA<br>11-118 | mA<br>11-204 | mA<br>11-205 | Aduca<br>numab | Bapineu<br>zumab | Gantene<br>rumab | Crenezu<br>mab | Donane<br>mab | Ponezu<br>mab | Solanez<br>umab |
| --- | --- | --- | --- | --- | --- | --- | --- | --- | --- | --- | --- |
| 6 | 209 | 133 | 13 | 71 | 18 | 164 | 241 | 355 | 1 | 276 | 96 |
| 7 | 77 | 61 | 8 | 37 | 4 | 66 | 101 | 259 |  | 136 | 32 |
| 8 | 46 | 15 | 1 | 24 | 2 | 32 | 51 | 155 |  | 65 | 11 |
| 9 | 21 | 9 | 0 | 8 | 1 | 17 | 10 | 85 |  | 30 | 3 |
| ≥10 | 41 | 24 | 1 | 9 | 1 | 26 | 18 | 142 |  | 28 | 4 |

**Table 3.**
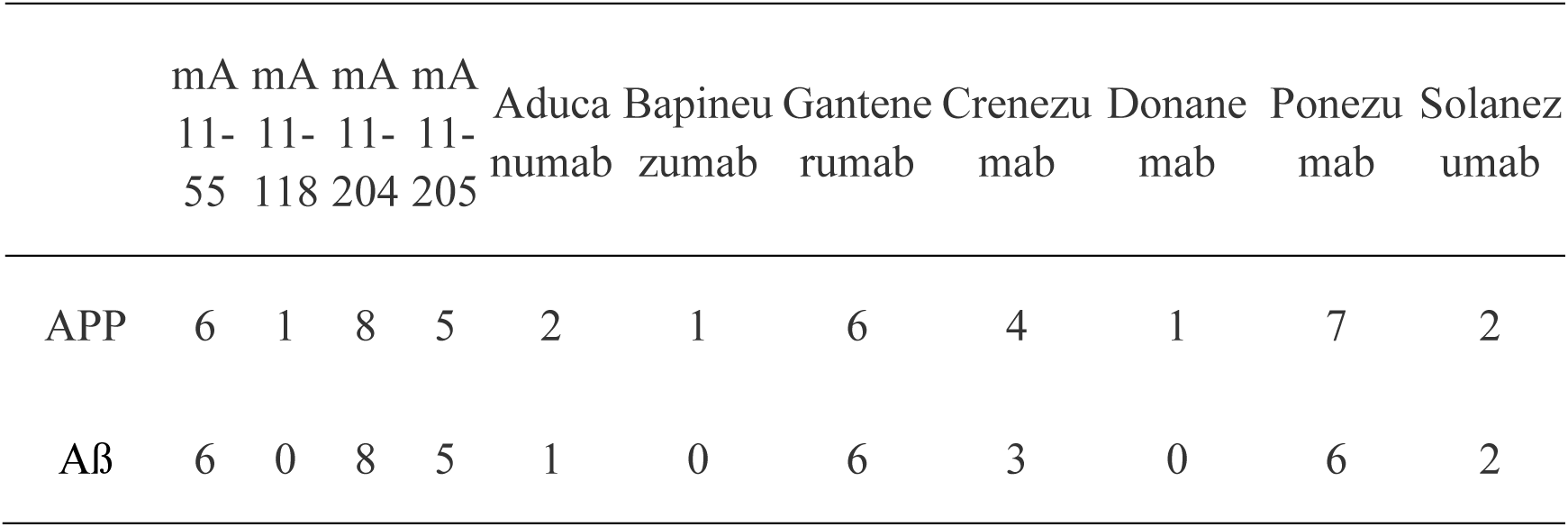
Statistics of strong interaction fragments between APP/Aβ and antibodies.

To investigate cross-reactivity between antibodies and proteins that form pathogenic or functional amyloid fibrils, a total of 74 Homo sapiens proteins were identified from the amyloid fibril-forming proteins database[66] (Supplementary information3). Each protein was divided into dodecapeptides based on the original sequence order, which were then used to predict interactions with antibodies using the best model. Peptides that had strong binding affinity with the antibodies were further analyzed. The non-specificity index of each antibody was calculated as the sum of the strongly bound peptides, which served as a measure of antibody specificity.

We investigate the binding possibilities between antibodies and proteins that are known to form amyloid fibrils from the database of amyloid fibril-forming proteins[66] (Supplementary information3). It can be seen from the figure 4(a)(b) that experimental antibodies typically exhibit higher specificity than drug antibodies, which raises thought-provoking questions. Fibrinogen α Chain antigen is most likely to cross react with antibody because it has the most twelve peptides that can strongly bind with antibody. Gastric inhibitory polypeptide antigen contains the least twelve peptides that can react with antibodies. As can be seen from Figure 4 (c), the non-specificity index of Crenezumab was the highest, reaching 5511. In contrast, Donanemab’s non-specificity index was the lowest, suggesting a lower probability of cross-reactivity. Notably, Aducanumab, which has been approved by the FDA for the treatment of Alzheimer’s disease[67], displayed better specificity compared to most other antibodies tested in the study.

**Figure 4.**
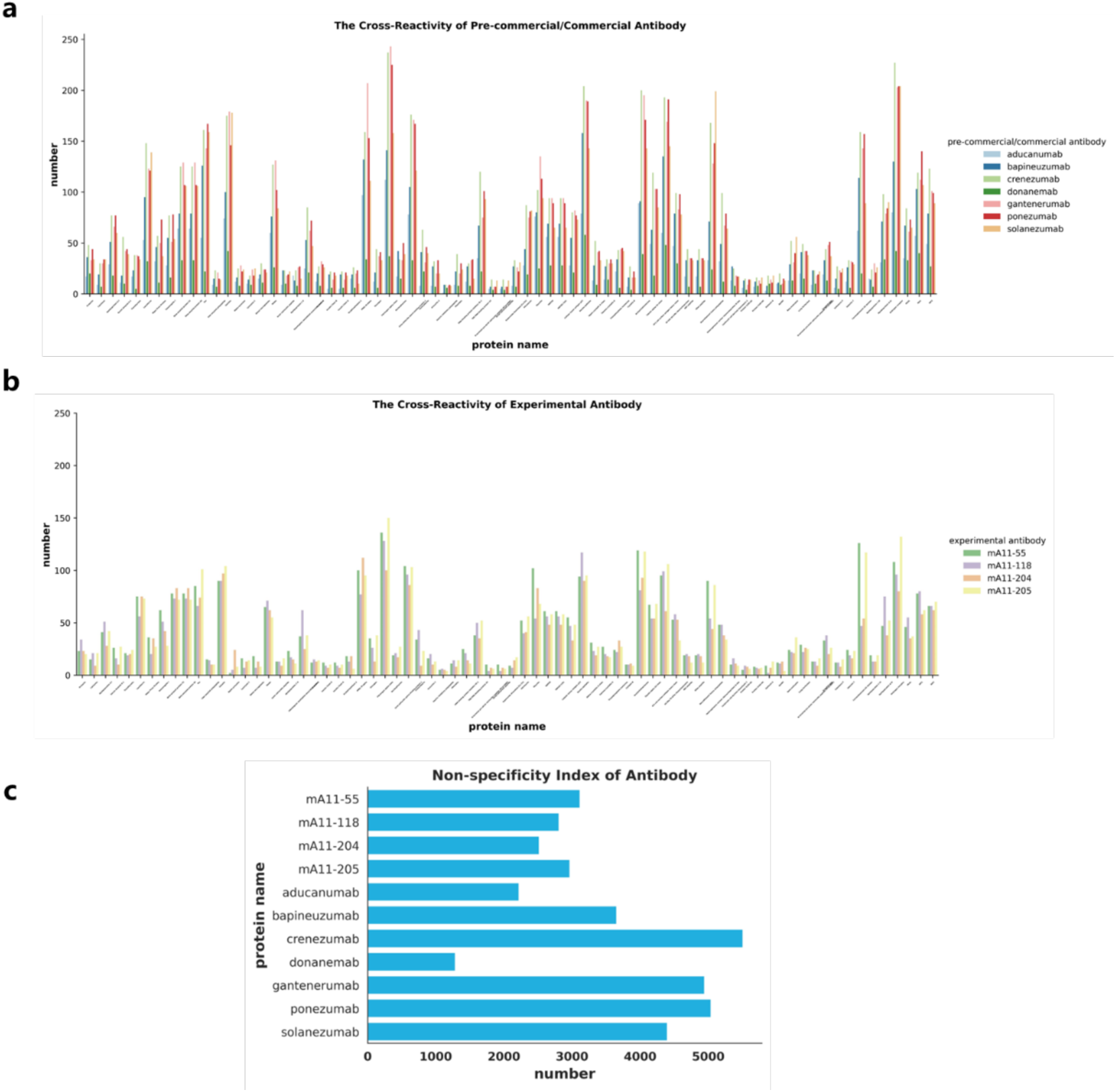
Predicting Antibody Specificity and Cross-Reactivity for Effective Amyloid-Targeted Therapeutics (a) Cross-reaction between commercial research antibody and protein that has been proved to form pathogenic or functional amyloid fibril (b) Cross-reaction between experimental antibody and protein that has been proved to form pathogenic or functional amyloid fibril (c) Non- specificity index of antibody

ABTrans was applied to study the off-target effects of antibodies. The sequence of 23391 human protein structures predicted by Aphafold[56] was segmented into dodecapeptide and then input into the model with 11 antibodies (4 antibodies in the experimental data[47] and 7 other commercial research antibodies[57–63]) to predict their possible interactions. Among these 11 antuibodies, Donanemab has the least strongly binding off-target protein. It could be explained by Donanemab recognizing AβpE3-42 a pyroglutamate form of Aβ that is aggregated in amyloid plaques [62], but our model did not learn pyroglutamate pattern from the data. Aducanumab is the first monoclonal antibody approved for the treatment of Alzheimer’s disease[67]. It can be seen from table 2 that Aducanumab interacts with fewer human proteins than other antibodies. The number of off-target proteins, binding site, and binding sequence of each protein were reported in Supplementary information3, hoping to guide the development of similar antibodies in the future.

The study further highlighted that Aβ antibodies may cross-react to multiple antigens, leading to a potential loss of specificity. This finding provides crucial insights into the challenges associated with the development of antibody-based therapeutics for Alzheimer’s disease and other amyloid-related disorders. Moreover, the study’s results may facilitate the design and selection of antibodies with higher specificity and efficacy, ultimately improving the outcomes of clinical trials and treatments for AD.

## Discussion

Antibodies against amyloid beta are promising therapeutics for Alzheimer’s disease. Predicting the interaction of antibodies and Aβ peptide is crucial for the development of more effective therapeutic agents. We have developed a deep learning model called ABTrans to predict the interaction between anti-Aβ antibodies and Aβ- like peptides. The AI model was trained based on the 1.5 million peptides screened from phage display experiment data[47] and 110 publicly known anti-Aβ antibody sequences. While original phage displays were screened using 28 anti-Aβ antibodies, the phage display identified peptides cover most Aβ1-42 region, with a distribution similarly observed when surveying known anti-Aβ antibody complex structures[41]. Therefore, it is a valid assumption that we can expand anti-Aβ antibody dataset to include all anti-Aβ antibodies with sequences available publicly.

Compared with common binary protein-protein interaction prediction, ABTrans can predict the affinity in four levels for the first time: strong, medium, weak, and non-binding. The accuracy of our model reached 0.83. The interpretability of attention is used to explain the interaction between Aβ and antibodies, and the results are consistent with that most antibodies recognize the N-terminal of Aβ peptide. ABTrans is applied to examine possible off-target binding of anti-Aβ antibodies, and we found that Aducanumab and Donanemab are indeed have the least off-target scores among the 11 antibodies screened in this work.

There have been many studies to predict antibody-antigen interaction using deep learning, such as predicting the paratope of antibody [68–71] and predicting the docking posture rank of antibody-antigen complex[72]. It should be noted that our model only considered linear peptides when screening human proteins. Prediction of conformational epitopes is more difficult than linear ones, [73] and the possible conformational epitopes from human proteome are extremely huge. However, while the anti-Aβ antibody binding peptides identified from phage display covers most of Aβ1-42 sequences, many of these peptides corresponding to conformational epitopes that anti-Aβ antibodies might recognize. Thus, the linear epitopes we have screened may also related to the conformational epitopes of anti-Aβ antibodies. Nevertheless, we will develop efficient algorithm to examine possible off-target bindings to conformational epitope in future study.

## Acknowledgments

1. B. Ma thanks support from Natural Science Foundation of China (Grant No. 32171246) and Shanghai Municipal Government Science Innovation grant 21JC1403700. Y. Wang thanks support from the grants from the National Natural Science Foundation of China (No.32200531), the Joint Research Funds for Medical and Engineering and Scientific Research at Shanghai Jiao Tong University (YG2022QN114 and YG2022QN082), and Startup Fund for Young Faculty at SJTU (SFYF at SJTU). All computation was performed using high performance computer cluster of Shanghai Jiao Tong University. We thank Ms. Y. Ge for helping collecting Aβ antibody sequences.

## Notes

### Competing Interest Statement

The authors have declared no competing interest.

